# pH-dependent 11° F_1_F_o_ ATP synthase sub-steps reveal insight into the F_o_ torque generating mechanism

**DOI:** 10.1101/2021.05.16.444358

**Authors:** Seiga Yanagisawa, Wayne D. Frasch

## Abstract

Most cellular ATP is made by rotary F_1_F_o_ ATP synthases using proton translocation-generated clockwise torque on the F_o_ c-ring rotor, while F_1_-ATP hydrolysis can force anticlockwise rotation and proton pumping. Although the interface of stator subunit-a containing the transmembrane half-channels and the c-ring is known from recent F_1_F_o_ structures, the torque generating mechanism remains elusive. Here, single-molecule studies reveal pH-dependent 11° rotational sub-steps in the ATP synthase direction of the E. coli c_10_-ring of F_1_F_o_ against the force of F_1_-ATPase-dependent rotation that result from H^+^ transfer events from F_o_ subunit-a groups with a low pKa to one c-subunit of the c-ring, and from an adjacent c-subunit to stator groups with a high pKa. Mutations of subunit-a residues in the proton translocation channels alter these pKa values, and the ability of synthase substeps to occur. Alternating 11° and 25° sub-steps then result in sustained ATP synthase rotation of the c_10_-ring.

## Introduction

The F_1_F_o_ ATP synthase (**Fig 1**) that is found in all animals, plants, and eubacteria is comprised of two molecular motors that are attached by their rotors, and by their stators (1,2). The Fo motor, which is embedded in bioenergetic membranes, uses a nonequilibrium transmembrane chemiosmotic proton gradient also known as a proton-motive force (pmf) to power clockwise (CW) rotation of its ring of c-subunits relative to the stator proteins as viewed from the *E. coli* periplasm. The c-ring docks to subunits-γand ε of F_1_. Subunit-γ, which serves as the drive shaft for F_1_, penetrates into the core of the F_1_ (αβ)_3_-subunit ring (**Fig 1B**) where each αβ-heterodimer comprises a catalytic site that synthesizes ATP from ADP and Pi.

**Fig. 1.**
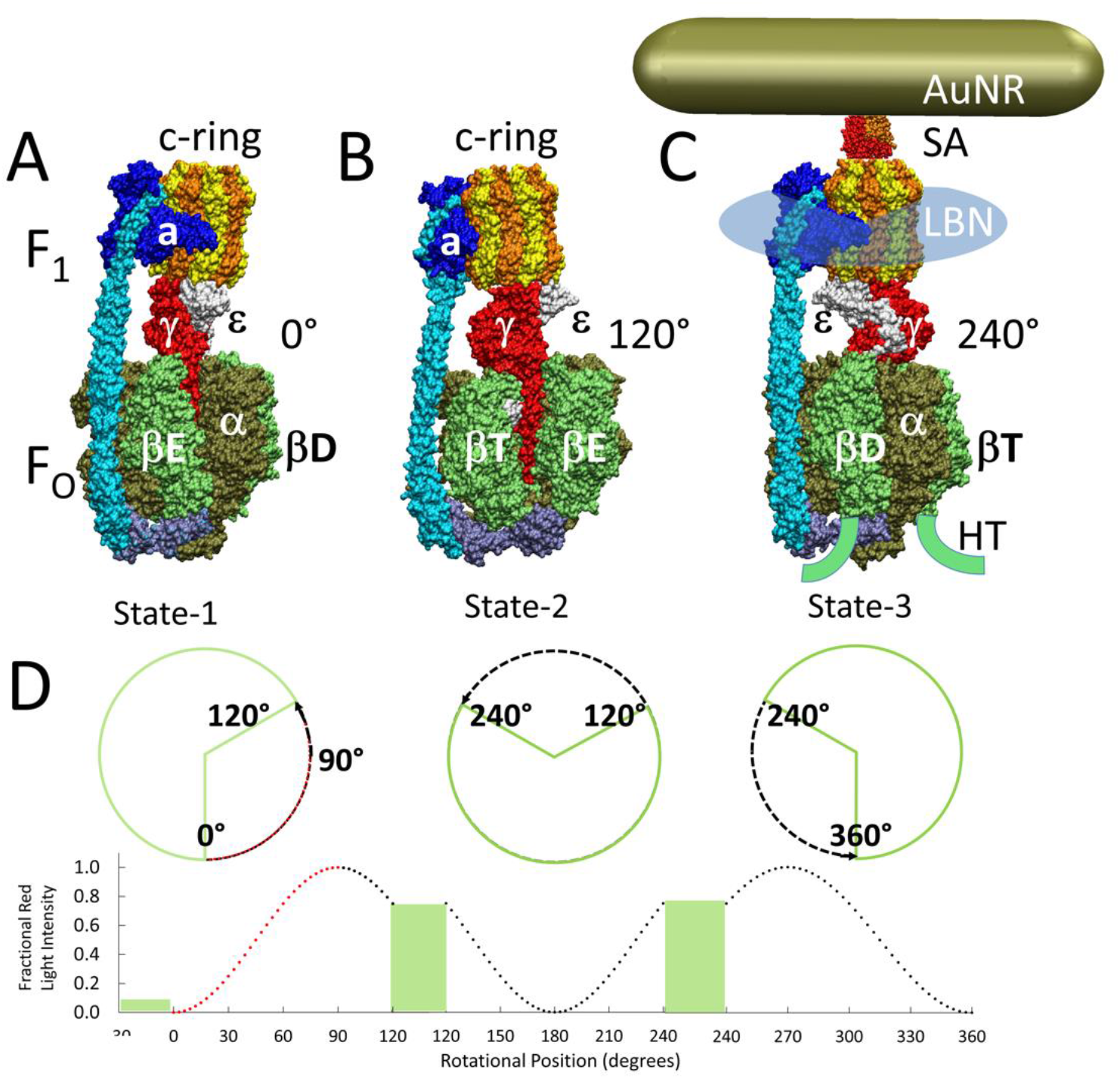
Cryo-EM structures of F_1_F_o_ ATP synthase inhibited by ADP in three rotary states, and measurement of changes in rotational position between catalytic dwells. **A** Rotational State-1, pdb-ID 6OQU (17). **B** State-2, pdb-ID 6OQV, with rotor 120° CCW from **A** where subunit-α is not shown to reveal subunit-γ. **C** State-3, pdb-ID 6WNR, with rotor 240° CCW from **A** showing microscope slide assembly of nanodisc-embedded F_1_F_o_ for rotation measurements. His_6_-tags on β-subunit C-termini enabled attachment to slide, while the streptavidin-coated gold nanorod (AuNR) bound to the biotinylated subunit-c ring. **D** Rotational position of single F1FO molecules versus time was monitored by intensity changes of polarized red light scattered from the AuNR in the presence of 1 mM Mg^2+^ ATP, which enabled F1-ATPase-dependent 120° CCW power strokes between catalytic dwells. Prior to data collection at 200 kHz, a polarizer in the scattered light path was rotated to minimize intensity during one of the three catalytic dwells. Light intensity increased to a maximum upon rotation by 90° during the subsequent CCW 120° power stroke. For each molecule the angular dependence of these power strokes versus time was analyzed.

Due to the staggered conformations of the three F_1_ catalytic sites, Site-1 contains ADP and Pi, and Site-2 contains ATP. Pmf-powered CW rotation of subunit-γforces conformational changes to all catalytic sites in the (αβ)_3_-ring, which releases ATP from Site-3 to create an empty site with each 120° rotational step (1,2). In this manner, F_1_F_o_ converts the energy from the pmf into a nonequilibrium chemical gradient (ΔμATP) where the ATP/ADP•Pi concentration ratio is far in excess of that found at equilibrium. The F_1_-ATPase motor can also use the energy from a ΔμATP to overpower the F_o_ motor and drive ATPase-dependent counter clockwise (CCW) rotation in 120° power strokes. Power strokes are separated by catalytic dwells, during which ATP is hydrolyzed (3–5). This rotation is used by F_o_ to pump protons across the membrane.

The means by which H^+^ translocation generates rotational torque on the c-ring is poorly understood. Maximal ATP synthase rates catalyzed by *E. coli* F_1_F_o_ ATP synthase are typically achieved with inner and outer pH values of 5.0 and 8.5, respectively, as measured using inverted membranes (6,7). Protons enter and exit F_o_ via half-channels in stator component subunit-a, which in *E. coli* F_1_F_o_, are separated by conserved arginine aR210 (1,2). During ATP synthesis, the input and output half-channels protonate and deprotonate, respectively, the carboxyl sidechain of conserved residue cD61 on each successive c-subunit in the *E. coli* c_10_-ring such that each H^+^ translocated results in a 36° rotation event.

Single-molecule studies revealed that when F_1_F_o_ is embedded in lipid bilayer nanodiscs (**Fig 1C**), the 120° CCW power strokes catalyzed by ATP hydrolysis were interrupted by transient dwells (TDs) at ~36° intervals corresponding to successive interactions between subunit-a and the c_10_-ring. In more than 70% of TDs, the F_o_ motor not only halted F_1_-ATPase CCW rotation, but the c-ring was able to rotate in the CW (ATP synthesis) direction. The occurrence of TDs increased inversely with pH between pH 5 and pH 7 (8), or when viscous drag on the nanorod was sufficient to slow the angular velocity of the F_1_-ATPase-driven power stroke (9,10).

Mutations of subunit-a residues aN214, aE219, aH245, aQ252, and aE196 of *E. coli* F_1_F_o_ decreased ATP synthase activity, and ATPase-dependent H^+^ pumping, supporting their participation in H^+^ translocation (10–13). Formation of TDs observed in single-molecule studies under viscosity-limited conditions also decreased significantly as the result of aE196 mutations (10). Although recent cryo-EM structures of F_1_F_o_ show that these residues are positioned along possible channels (2,14–17), some are not protonatable, and most are separated by distances too large to form hydrogen bonds.

We have now determined the pKa values of TDs in single-molecules of F_1_F_o_ embedded in a lipid bilayer nanodiscs. These studies reveal that the CW rotation TDs occurs in pH-dependent 11° ATP synthase sub-steps that depend on H^+^ transfer between protonated groups with a low pKa from the subunit-a input channel to the c-ring, and between the c-ring to unprotonated groups with a high pKa in subunit-a. Mutations of residues that participate in H^+^ translocation in the input and output channels change both pKa values, and alter the probability of forming 11° ATP synthase sub-steps. These data support a mechanism where sustained c-ring rotation in the ATP synthesis direction results from successive alternating 11° and 25° sub-steps for each c-subunit in the c_10_-ring.

## Results

Contributions of subunit-a residues putatively involved in the ATP synthase H^+^ half-channels were assessed by the effects on transient dwell formation caused by mutations that converted charged or polar groups in subunit-a to hydrophobic leucine. Changes in rotational position were measured by a 35 x 75 nm gold nanorod (AuNR) bound to the biotinylated c-ring of individual *E. coli* F_o_F_1_ molecules embedded in lipid bilayer nanodiscs (9), hereafter F_1_F_o_ (**Fig 1C**). Changes in rotational position during F_1_-ATPase power strokes in the presence of saturating 1 mM MgATP were monitored by the intensity of polarized red light scattered from the AuNR (18,19). Prior to data collection, the polarizer was adjusted so that the scattered red light intensity was at a minimum during one of the three F_1_ catalytic dwells (**Figs 1D, 2A**). The subsequent power stroke caused an increase in light intensity to a maximum when the AuNR had rotated 90° (20). Rotational data sets of each F_1_F_o_ molecule examined were collected for 5 sec, which included ~300 of these power strokes (8). Ten data sets were collected for each molecule. The number of F_1_F_o_ molecules examined at each pH for WT and mutants is indicated in Supplementary Figure 3. Using WT at pH 5.0 as an example where data from 103 F_1_F_o_ molecules were collected, this was equivalent to 1030 data sets, and ~309,000 power strokes examined. For each molecule examined, rotational position versus time was calculated from scattered light intensity versus time using an arcsine^1/2^ function from which the number of TDs observed during the first 90° of rotation were determined (21).

**Fig. 2.**
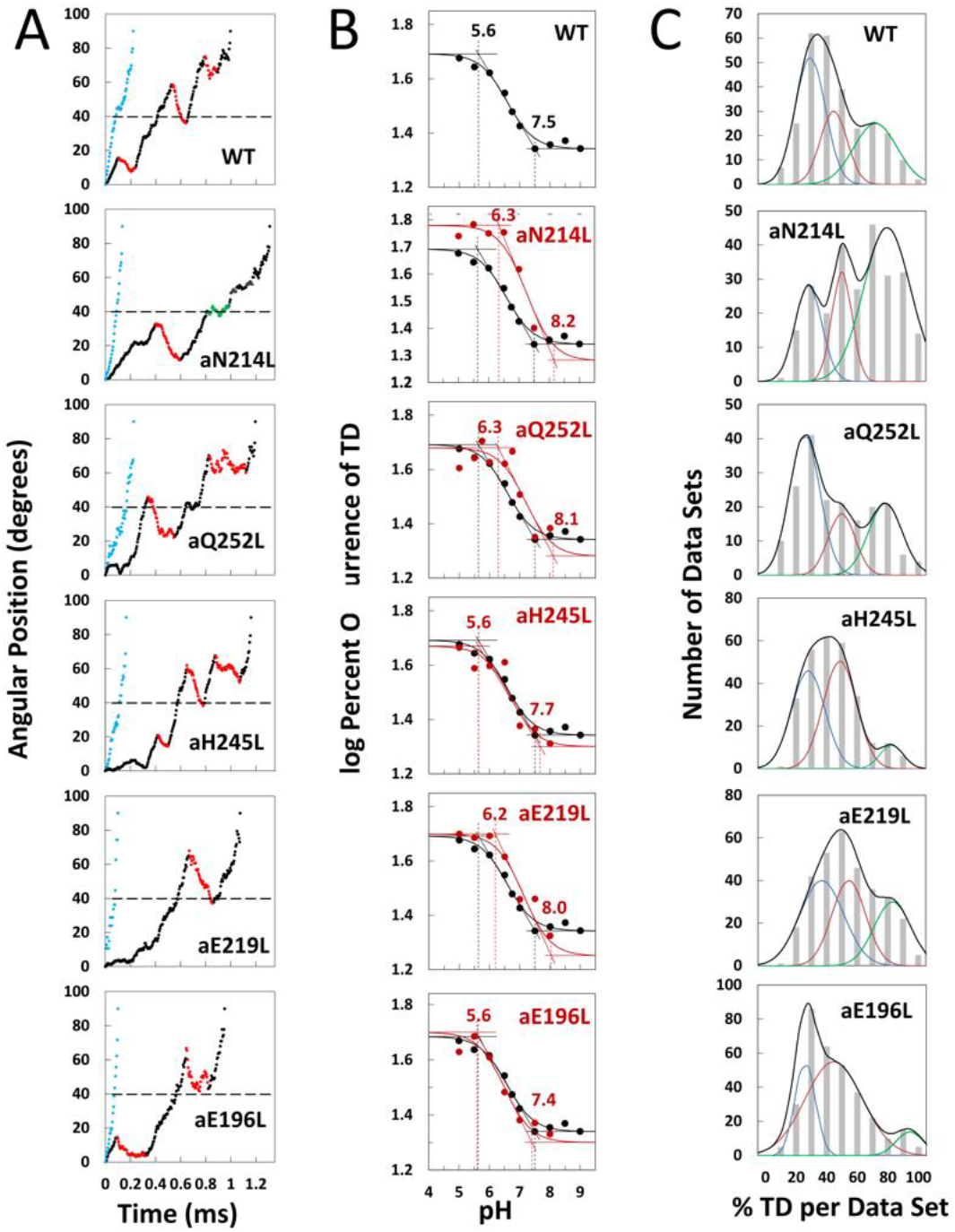
Effects of subunit-a mutations on Transient Dwells (TDs). **A** Examples of power strokes without transient dwells 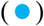, and of power strokes with transient dwells that lacked 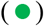, or contained CW c-ring rotation relative to subunit-a 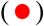 plotted as degrees of rotation after the catalytic dwell *vs* time where 40° 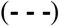 is the optimal position for binding of ATP or inhibitory ADP (3, 28). **B** Average percent TDs per data set *vs* pH from which pKa values were derived via intercepts of the slope and plateaus 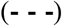 of each curve based on the fit of the data to *Eq. 1* for WT 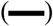, and subunit-a mutants 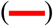. Distribution of the extent of synthase step CW rotation at pH values when the percent of synthase steps was minimum 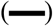, and maximum 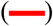. **C** Distributions at pH 6.0 of the percent of TDs per data set of power strokes (gray bar graphs) where multiple data sets that each Slide2contained ~300 power strokes were collected from each of the total number of the F_1_F_o_ indicated, and data were binned in 10 % increments. The data were fit to the sum of three Gaussians 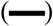 representing low 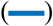, medium 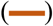, and high 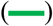 efficiencies of TD formation.

Examples power strokes from WT and mutant F_1_F_o_ molecules at pH 5.0 where TDs were present (•), and absent 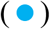 are shown in Fig 2A and Fig S1. When present, TDs either stopped F_1_-ATPase CCW rotation momentarily 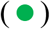, or exhibited CW rotation in the ATP synthase direction, hereafter synthase steps 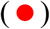. None of the mutations examined eliminated the ability of F_1_F_o_ to form TDs. Power strokes typically contained 2 to 3 TDs, when present. These were separated by an average of ~36°, consistent with an interaction between subunit-a and successive c-subunits in the c_10_-ring of *E. coli* F_1_F_o_.

### Subunit-a Mutations Alter pKa’s of TD Formation

We determined the pKa values of groups that contribute to TD formation (**Fig 2B**) using equations applied to the pH dependence of enzyme inhibition kinetics (22). Transient dwells occur when subunit-a binds to the c-ring to stop F_1_ ATPase-driven rotation for a period of time. Thus, a TD represents an extent that F_o_ inhibited the F_1_ ATPase motor, which occur as often as 3.6 times per F1 power stroke. Kinetically, the ATPase power stroke duration without TDs is ~200 μsec, while TDs each last ~100 μsec (8,9). In data sets where TDs occur in 100% of the power strokes, e.g. aN214L at pH 6.0, this represents a 64% inhibition of the F_1_ATPase power stroke kinetics.

A maximum average of 47.5% of WT power strokes from all three efficiency groups occurred at pH 5.0, which decreased with increasing pH until it plateaued at a minimum of ~22% at pH values >7.5 (**Fig 2B**). The pH dependences for WT and mutants were fit to *Eq. 1* where T is the total average TD occurrence, T_min_ is the minimum TD occurrence, and K_1_ and K_2_ are the inhibition constants that define the increase and maximum TD occurrence versus pH as the result of either a residue that is protonated with pKa_1_, or unprotonated with pKa_2_, respectively. It is noteworthy that K_1_ is similar to a dissociation constant because a smaller K_1_ increases the ability of subunit-a to bind to, and stop c-ring rotation with decreasing pH (**Fig S2**). Conversely, a smaller K_2_ value decreases TD formation with decreasing pH because it is the unprotonated form of that residue that binds and inhibits.

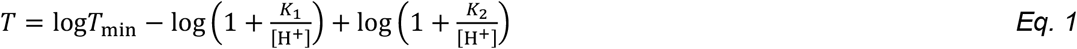

The fit of the data to *Eq. 1* defines the slope of the curve as well as the high and low plateau values. Because these are log-log plots, the pKa values (**Fig 2B**, 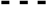) are determined by the intercept of the slope with the high and low plateau values 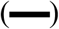. None of the mutations changed T_min_ significantly. Using parameters derived by the fits of the data to *Eq. 1* for WT and mutants (**Table 1**), the WT group(s) that must be protonated to induce a TD had pKa_1_ and K_1_ of 5.6, and 6.4, respectively, while the group(s) that must be unprotonated to induce a TD had pKa_2_ and K_2_ values of 7.5 and 6.75.

**Table 1.** pKa values and inhibition constants for WT and subunit-a mutants. Values were derived from the fits to *Eq. 1* of the average percent of TDs per data set *vs* pH in Fig 2C.

| | $K_1$ | $K_2$ | $T_{\min}$ (%) | pKa<sub>1</sub> | pKa<sub>2</sub> |
| --- | --- | --- | --- | --- | --- |
| WT | 6.4 | 6.75 | 22.0 | 5.6 | 7.5 |
| aN214L | 7.0 | 7.50 | 19.1 | 6.3 | 8.2 |
| aQ252L | 7.0 | 7.40 | 19.5 | 6.3 | 8.1 |
| aE219L | 6.9 | 7.35 | 17.8 | 6.2 | 8.0 |
| aH245L | 6.5 | 6.87 | 20.0 | 5.6 | 7.7 |
| aE196L | 6.3 | 6.70 | 20.0 | 5.6 | 7.4 |

The aN214L mutation, which had the greatest effect on the pH dependence of TD formation, increased the maximum percent of TDs formed at low pH to 61% (1.3-fold), and shifted the pH dependence in the alkaline direction from WT. These changes were due to increases in K1 and K2 to 6.4 and 6.75, respectively, that increased pKa_1_ and pKa_2_ by 0.9 and 0.7 pH units. The differential increases in K_1_ and K_2_ by 0.6 and 0.75 units led to the aN214L-dependent increase in maximum TD formation at low pH because an equal shift of these values in the same direction causes the curve to shift to higher pH values without affecting the maximum occurrence of TDs formed (**Fig S2**). Similar but smaller effects were observed with aQ252L, and aE219L (**Fig 2B**) where K_1_ increased by 0.6 and 0.5 units, respectively, resulting in an pKa_1_ increase of almost 1 pH unit from that of WT. However, aQ252L, and aE219L decreased K_2_ by 0.35 and 0.40 units from WT such that the increase in pKa2 was proportionally smaller than that observed for aN214L. Consequently, while both mutants shifted the pH dependence in the alkaline direction from that of WT, only aQ252L showed an increase in the maximum TD occurrence (52%).

Mutations aH245L and aE196L caused the smallest changes on the pH dependence of TD formation. The former increased K_1_ and K_2_ by 0.1 and 0.12 units, which had no effect on pKa_1_, and increased pKa_2_ by 0.2 units. The latter was the only mutation to decrease the values of both K_1_ and K_2_, which decreased pKa_2_ by 0.1 pH units from that of WT. It is noteworthy that aE196 is a component of the H^+^ output channel.

### Subunit-a Mutations Affect TD Formation Efficiency

The percent of TDs observed per data set fit to three Gaussian distributions with low 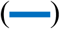, medium 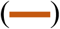, and high 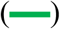 efficiencies as shown at pH 6.0 (**Fig 2C**), and at all pH values examined (**Fig S3**). Subunit-a mutations affected the percent of TDs formed per data set during power strokes in each of these efficiencies, which correlate to the three rotary positions of the central stalk relative to the peripheral stalk (8). The proportional differences of efficiencies of TD formation is shown relative to the average low efficiency for WT (**Fig 3**). Medium and high efficiency distributions of TDs in WT increased 1.5-fold and 2.2-fold, respectively, relative to low efficiency. The aN214L mutation increased the percent of TDs per data set for high, medium, and low efficiencies by 3-fold, 2-fold, and 1.2-fold, respectively, from the WT low efficiency. Mutations aQ252L and aE219L also increased TDs per data set for the high (2.7-fold and 2.5-fold), and medium (1.7-fold and 1.6-fold), but not the low efficiency distributions. Mutations aH245L and aE196L either did not increase the efficiency or slightly decreased the efficiency of the distributions of TD formation per data set.

**Fig. 3.**
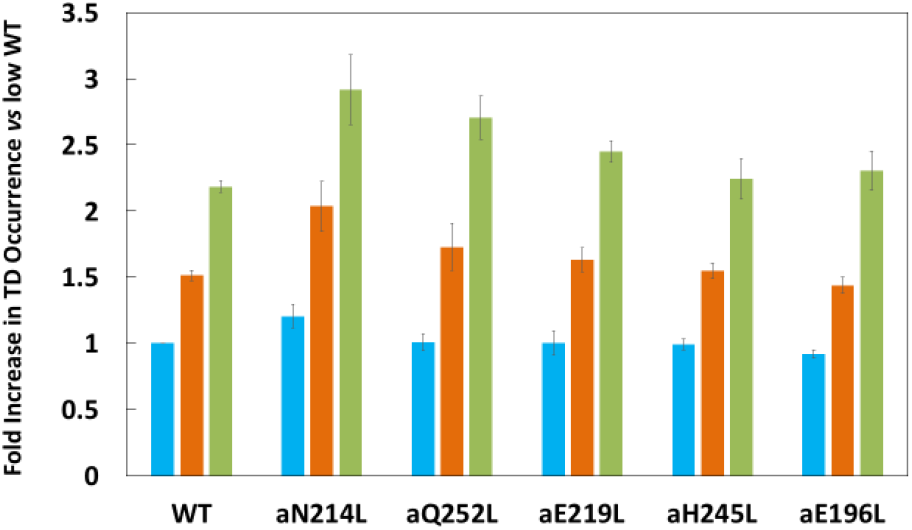
Proportion of low 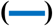, medium 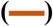, and high 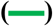 TD formation efficiencies relative to WT low efficiency TD formation. Each was the average of all pH values examined. Vertical bars represent standard error.

### Synthase Steps Rotate CW an Average of ~11°

The proportion of TDs with and without a synthase step for WT and mutants are shown in **Fig 4A** at the pH values when the proportion of synthase steps was minimum 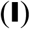 and maximum 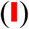, and at all pH values examined in Fig S4. The minimum proportion of synthase steps was observed at pH 5.5 for WT and all mutants except aN214L that occurred at pH 6.0. Even at this low pH values, synthase steps accounted for 62% - 68% of all TDs. In WT, a maximum of ~80% of TDs contained synthase steps at pH 7.0, which was an increase of 13% from the minimum. These plots also show the distributions of the extent of CW rotation during a synthase step, for which the 11° and 9° average and median values of CW rotation, respectively, were not changed significantly by the mutations (**Fig 4B**).

**Fig. 4.**
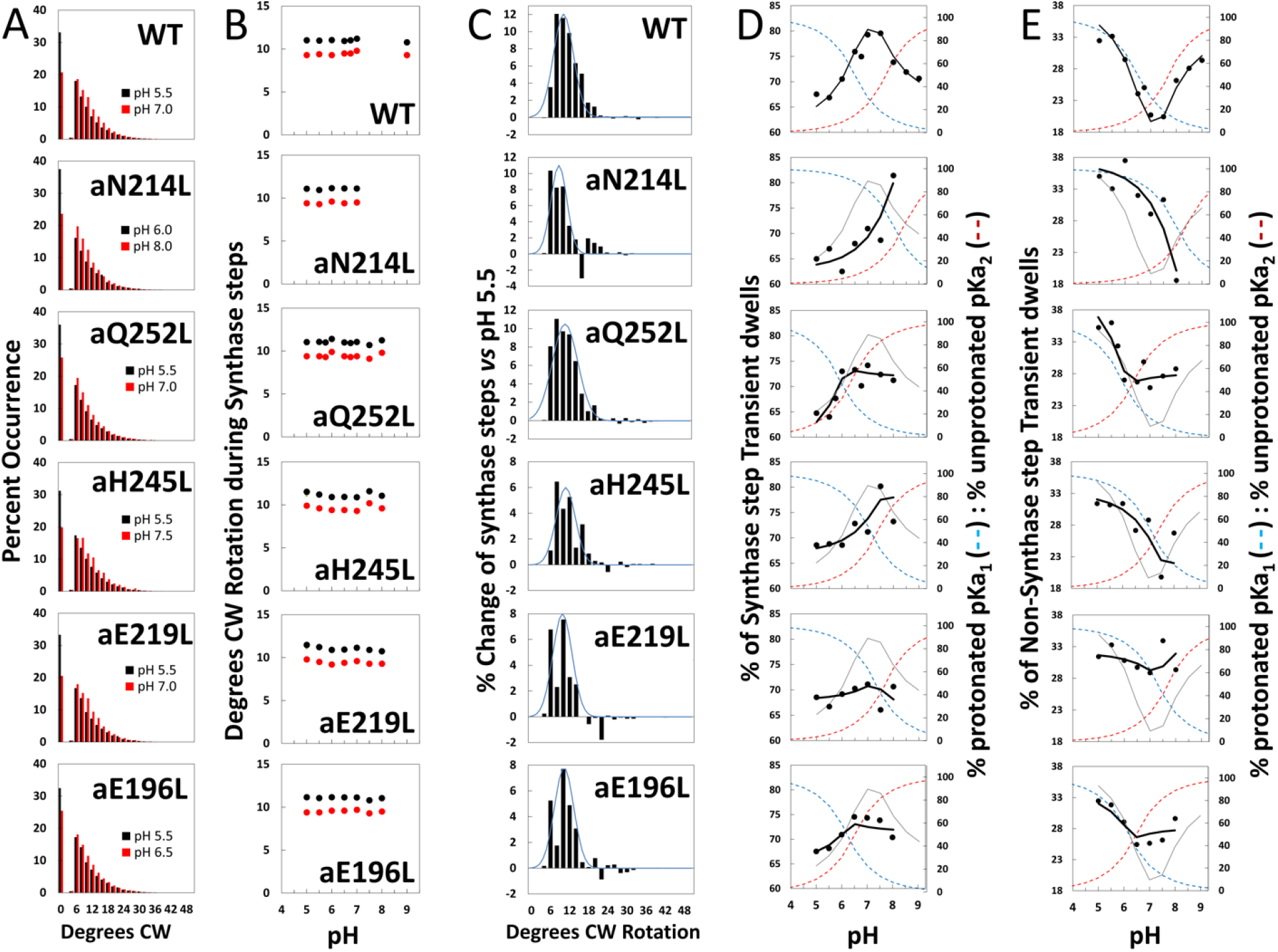
Effects of subunit-a mutations on the pH dependence of the extent of CW synthase step rotation, and fraction of TDs containing synthase steps. **A** Distributions of the extent of CW rotation in the ATP synthesis direction during transient dwells for WT and subunit-a mutants at the low (black) and high (red) pH values indicated. **B** Mean (•) and median 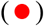 extents of CW rotation during a synthase step *vs* pH. **C** Distributions of the difference in extent of CW synthase step rotation between pH values in Fig 2D when the percent of synthase steps was maximum vs. minimum where 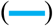 is the Gaussian fit. **D** Percent of TDs containing CW synthase steps *vs* pH, where the data were fit to *Eq. 3* 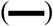. The fraction of protonated groups with pKa_1_ 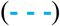, and unprotonated groups with pKa_2_ 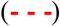 *vs* pH was calculated from the pKa values of Table 2. **E** Percent of TDs that lack synthase steps *vs* pH where the probability of forming a TD without a synthase step 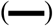 was determined by *Eq. 2* from the fraction of protonated groups with pKa_1_ 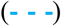, and unprotonated groups with pKa_2_ 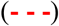 *vs* pH calculated using pKa values from Table 2.

After subtracting the occurrence of the extent of synthase step CW rotation at the pH when it was at a minimum 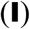 from that observed at other pH values (**Fig S4**) including that at its maximum 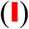, a Gaussian distribution of the increase in the extent of synthase step CW rotation was observed (**Fig 4C**). During a synthase step, the mean and standard deviations in the extent of CW rotation was 12°±3° for WT, with little variation resulting from the mutations including: 11°±3° (aN214L), 11°±4° (aQ252L), 11°±3° (aH245L), 10°±3° (aE219L), and 11°±3° (aE196L). In all cases, the distributions were truncated with minimum CW rotational steps of 6°. At their maxima, the extents of CW c-ring rotation during synthase events rotated 24° and 36° about 1% and 0.1% of the time, respectively.

### Subunit-a Mutations Affect the Proportion of TDs with Synthase Steps

The subset of TDs that forced the c-ring to rotate CW (synthase steps) against the CCW force of F_1_-ATPase rotation was pH dependent (**Fig 4D**). A maximum of 80% of TDs contained synthase steps in WT at ~pH 7.3, and a minimum of 67% at pH 5.5. At pH values >7.5, the proportion of synthase steps decreased to 71% at pH 9.0.

Because a TD either contains (T_S_) or lacks (T_N_) a synthase step, the pH dependence of TDs with a synthase step (**Fig 4D**) was the inverse of that without a synthase step (**Fig 4E**) per *Eq. 2*.

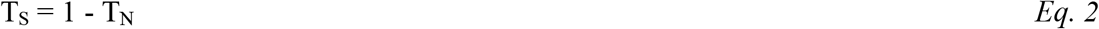

For WT, the minimum T_N_ of 20% at pH 7.5 increased 1.7-fold at pH 5.5, and also increase 1.5-fold and pH 9.0. At these extremes of low and high pH values, TD formation was dominated by groups where either pKa_1_ is protonated, or by unprotonated groups with pK_2_. This conclusion is supported by the good fits of the pH dependencies of TDs without synthase steps for WT and subunit-a mutants (**Fig 4E**) to *Eq. 3*, where the probability of forming a TD without a synthase step (T_N_) is the sum of the probability (P_1_) of the protonated group(s) with pKa_1_ (X_1_), and the probability (P_2_) of unprotonated group(s) with pKa_2_ (Y_2_). Thus, these results support the conclusion that a TD without a synthase step can result from a H^+^ transfer event from the protonated group with pKa_1_, *or* from a H^+^ transfer event to the unprotonated group with pKa_2_.

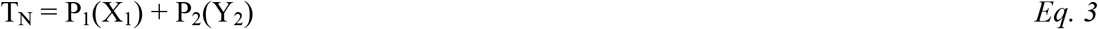

Fits of the pH dependence of TDs without synthase steps from *Eq. 3* 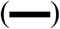 were based on the pKa values (**Fig 4E**), and probabilities summarized in **Table 2**. The WT data fit to probabilities of 38% and 33% for protonated groups (pKa_1_ 6.6), and unprotonated groups (pKa_2_ 7.7), respectively, such that the difference between the pKa values was 1.2 pH units. Consequently, T_N_ showed a minimum at ~pH 7.3, and maxima at high and low pH values when only the group(s) with either pKa_1_ or pKa_2_ were protonated and unprotonated, respectively.

**Table 2.** pKa values and probabilities of forming TDs without synthase steps for WT and subunit-a mutants. Values were derived from the fits of the data of Fig 4C to *Eq. 2*.

| | $pK_{a1}$ | $P_1$ (%) | $pK_{a2}$ | $P_2$ |
| --- | --- | --- | --- | --- |
| WT | 6.5 | 38 | 7.7 | 33 |
| aN214L | 8.0 | 37 | 8.4 | 5 |
| aQ252L | 5.9 | 42 | 6.4 | 28 |
| aE219L | 7.1 | 32 | 7.4 | 35 |
| aH245L | 7.3 | 33 | 7.7 | 22 |
| aE196L | 6.2 | 34 | 6.5 | 28 |

As a result of the subunit-a mutations, P_1_ values changed to a smaller extent (32% - 42%) than did P_2_ values (5%-35%). Except for aE219L, all mutations decreased P_2_, including a >6-fold decrease with aN214L. The difference between pKa values observed with the mutants was from 0.3 to 0.5 pH units compared to the 1.2 pH unit difference of WT. Both pKa_1_ and pKa_2_ of aN214L increased by 1.5 and 0.7 pH units such that the minimum T_N_ of ~18 % at pH 8.0 represented an increase of 0.7 pH units from that of WT. At pH 5.5, T_N_’s comprised 38% of all TDs in aN214L. A similar but smaller shift of the minimum T_N_ occurrence to pH 7.5 was also observed for aH245L, which primarily resulted from an increase in of pKa_1_ by 0.8 pH units from WT. A striking effect of mutations aQ252L, aE219L, and aE196L was that they suppressed the pH dependence of synthase step formation. Of these, aE219L was most where T_S_ varied between 66% and 71% of TDs over the pH range examined.

In all cases, the occurrence of synthase steps reached a maximum at the crossover point between the fractions of protonated groups with pKa_1_ and unprotonated groups with pKa_2_. This is the point at which the largest fractions of both groups were in the correct protonation state where H^+^ transfer events could occur from the pKa_1_ groups to the c-ring, *and* from the c-ring to the pKa2 groups.

## Discussion

The results presented here reveal that F_1_F_o_ generates pH dependent 11° CW rotational ATP synthase sub-steps of the c-ring relative to subunit-a that can occur against the force of F_1_ATPase-dependent CCW rotation at saturating ATP concentrations. Subunit-a residues most closely linked to these H^+^ transfer-dependent rotational steps are aN214/aQ252 in the input channel, and aE196/aS199 in the output channel based on results presented here that include: (i) that synthase steps, which occur in at least 67% of all TDs, are dependent on a group of residues with an average pKa of 6.5 that must be protonated, and a second group with an average pKa of 7.7 that must be unprotonated; (ii) the probability of forming an ATP synthase step reaches a maximum of 80% of TDs at pH 7.5 when the fractions of protonated groups and unprotonated groups with low and high pKa values are optimal; and (iii) mutating subunit-a residues in either the input or output half-channels alters both pKa values, and can decrease the fraction of TDs that exhibit synthase steps. Based on these results, we conclude (**Fig 5A → 5B**) that the 11° ATP synthase steps result from a H^+^ transfer event from the input channel residues aN214/aQ252 to the leading cD61 (pink), and a H^+^ transfer event from the lagging cD61 (orange) to output residues aE196/aS199.

**Fig. 5.**
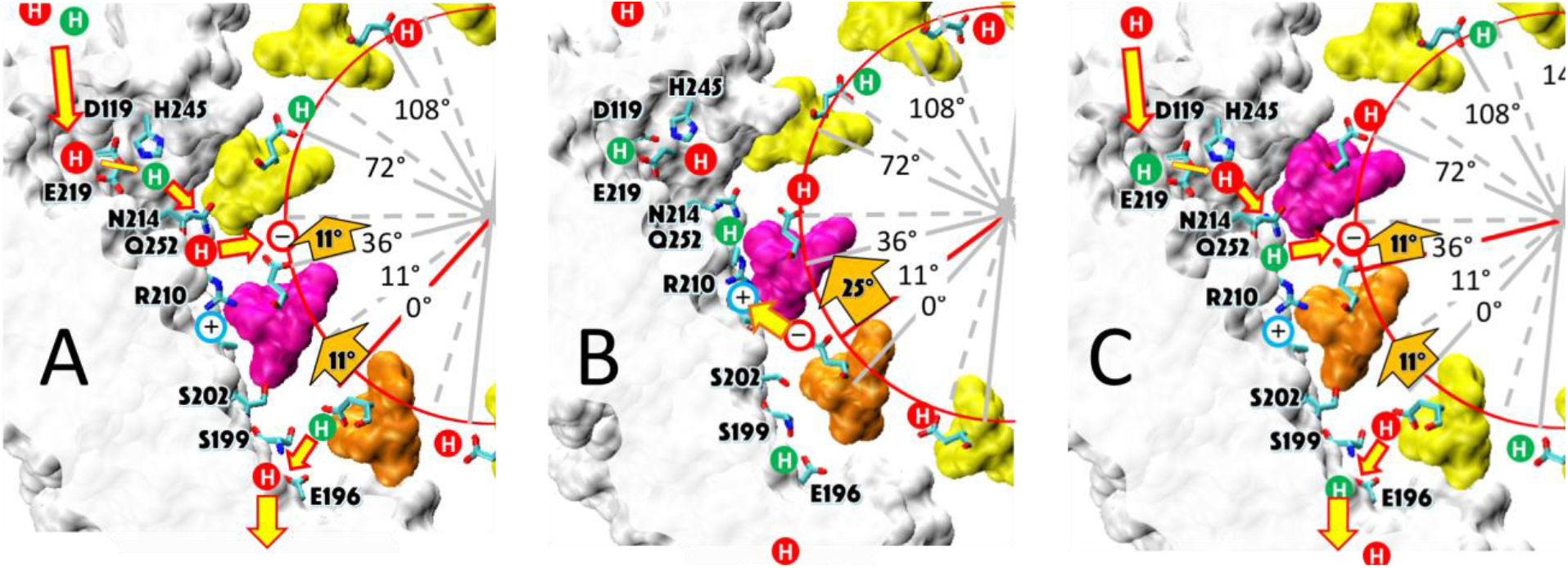
Alternating 11° and 25° sub-steps power F_o_ c-ring CW rotation in the ATP synthase direction. **A** The pH-dependent 11° sub-step occurs when H^+^ transfer from aN214/aQ25-bound water to the unprotonated leading cD61-carboxyl (pink), and from the protonated lagging cD61-carboxyl (orange) to aS199/aE196-bound water. Rotation results as the negatively charged lagging cD16 (orange) moves in response to the decrease in hydrophobicity from the lipid bilayer to that of the subunit-a interface. This decreases the distance between the lagging cD61 carboxyl and the aR210-guamdirnum from ~11.5 Å to ~7.5 Å. **B** The 25° sub-step occurs from the electrostatic interaction between the lagging cD61 carboxy (orange) and the aR210 guanidinium. **C** The electrostatic attraction decreases the distance between orange cD61 and aR210 from ~7.5 Å to ~3.5 Å to complete a 36° stepwise rotation of the c-ring, and positions the orange cD61 to become the leading carboxyl for the next pH-dependent 11° sub-step. The *E. coli* F_1_F_o_ cryo-EM structures of rotary sub-states pdb-IDs 5OQS (**A** and **C**), and 5OQR (**B**) are shown as cross-sections of subunit-a (white), and the c-ring as viewed from the periplasm.

A mechanism where F_o_ uses alternating 11° (**Fig 5A → 5B**), and 25° (**Fig 5B → 5C**) sub-steps to power c-ring rotation that drives ATP synthesis is consistent with the data presented here, and with F_1o_ structures. The pH-dependent 11° sub-step occurs upon H^+^ transfer from water bound to aN214 and aQ252 residues to the unprotonated leading cD61-carboxyl (pink), and from the protonated lagging cD61-carboxyl (orange) to the aS199 and aE196-bound water (**Fig 5A**). Rotation results as the negatively charged lagging cD61 moves from the lipid bilayer in response to the decrease in hydrophobicity at the interface with subunit-a, which contains polar groups above (aS202 and aS206) and below (aK203 and aY263) the plane of cD61 rotation (**Fig 5B**, and **Fig 6A**). The 11° sub-step decreases the distance between the lagging cD61 carboxyl and the aR210-guanidinium from ~11.5 Å to ~7.5 Å (**Fig 5B**), where we postulate that the electrostatic attraction between them becomes sufficient to induce the 25° sub-step. As the result of this sub-step, the distance between these groups decreases from 7.5 Å to 3.5 Å (**Fig 5C**). The loss of negative charge when the lagging cD61-carboxy is protonated by aN214 and aQ252 then allows this c-subunit to rotate away from aR210 into the lipid bilayer as the 11° sub-step repeats.

**Fig. 6.**
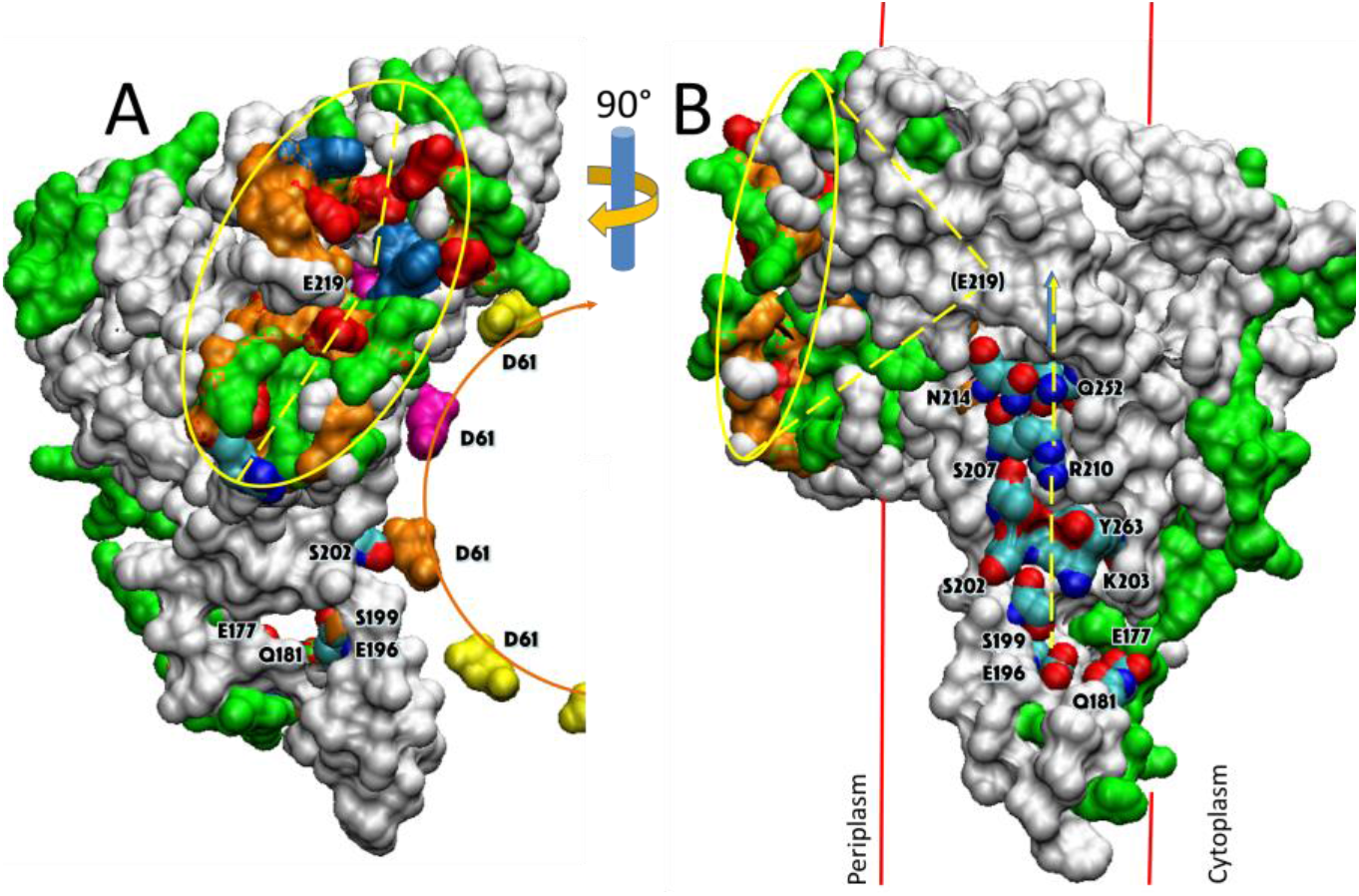
Aqueous vestibule of charged and polar groups can serve as an antenna to funnel protons to the input channel. **A** Periplasmic surface of subunits-a and b showing the aqueous vestibule (yellow oval) that can serve to funnel protons to aE219 (pink). The funnel surface is defined by asp and glu groups (red), his groups (blue), polar residues (green), and backbone residues of loop regions (orange). Hydrophobic residues are white. The cD61 carboxyl groups that interface subunit-a are indicated yellow, orange and pink. **B** Surface of subunits-a and b that interfaces the c-ring from structure pdb-ID 5OQR showing approximate position of the aqueous funnel terminating at aE219 (buried), and residues involved in proton translocation. The plane of cD61 residues and the direction of rotation in the ATP synthase direction is indicated by 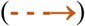.

The alternating 11° and 25° sub-step mechanism proposed here is consistent with *E. coli* F_1_F_o_ structures pdb-IDs 6OQR and 6OQS (17), which were used to illustrate Fig 5B and 5C, respectively. Although subunit-γ is docked to the leading c-subunit (pink) in both structures, the rotary position of the c-ring Fig 5C is 25° CW from that of Fig 5B. With a c_10_-ring, the cD61-carboxyls are positioned every 36°, consistent with the spacing between transient dwells observed here. Consequently, rotary positions of the cD61-carboxyls in Fig 5A and 5C are equivalent, except that we have labelled the leading cD61 in the former as the lagging cD61 in Fig 5C to demonstrate the 11° difference between Fig 5A and 5B.

Consistent with the pH-dependence of the 11° synthases steps observed here, the positions of the leading and lagging cD61-carboxyls to the subunit-a input and output residues in structure PDB-ID 5OQR (**Fig 5A**) suggest that they are poised to undergo H^+^ translocation events. Notably, the lagging cD61-carboxyl (orange) is ~3.5 Å from the aS199-hydroxyl, which is close enough to form a hydrogen bond to the protonated carboxyl group. The negatively charged leading cD61-carboxyl (pink) is ~3.8 Å from the aR210-guanidinium, which suggests that they have formed a salt bridge. However, the carboxyl is proximal to aN214/aQ25 where protonation would be most effective in allowing it to rotate CW. The FO conformation after H^+^ translocation is consistent with structure 6OQR (**Fig 5B**) where the c-ring has rotated 11° CW relative to subunit-a (17). Post H^+^ transfer, the aR210-guanidinium is now 5.1 Å away from the leading cD61 carboxyl (pink), which suggests that the carboxyl has been protonated. The lagging cD61-carboxyl (orange) is now 8.6 Å away from aS199, and its distance to the aR210-guanidinium has decreased from ~11.5 Å in Fig 5A to ~7.5 Å (**Fig 5B**). At this distance, and with the reduced polarity in the membrane, the electrostatic interaction between the unprotonated cD61 and the guanidinium group would be substantial. We postulate that this interaction is sufficient to power the 25° CW rotational sub-step to reset the conformation to that of structure PDB-ID 6OQS (**Fig 5C**).

The mechanism proposed here is also consistent with structures of the c-ring determined as a function of pH (23). These show the pH-dependent interconversion of the cD61 carboxyl between a protonated locked or closed conformation in subunit-c in a hydrophobic environment, and an unprotonated open, conformation in a more polar environment. Molecular dynamic simulations of these c-ring structures (23) found that it is energetically favorable for the unprotonated cD61 of the c-ring to form an ion pair with a nearby arginine bound to a peptide that was modeled in a lipid environment.

The rotational sub-state structures of *E. coli* F_1_F_o_ that differ by the 25° rotation of the c-ring relative to subunit-a were obtained when the complex was inhibited by ADP (17). Similar 11°, and 25° rotational sub-states have also been observed with ADP-inhibited F_1_F_0_ *B. taurus* (15), and *M. smegmatis* (24). In *M. smegmatis* F_1_F_o_, the binding of bedaquiline stabilizes a rotational sub-state that is either 25° CW, or 8° CCW from the equivalent rotational state in the absence of the drug (24). The rotational position of the c-ring in the cryo-EM structure of *S. cerevisiae* F_1_F_o_ is also changed by ~9° when the inhibitor oligomycin is bound to F_o_ (25).

The low, medium, and high efficiencies of TD formation reported here (**Fig 2B**) were attributed to torsional strain resulting from the asymmetry between 36° c_10_-ring stepping, and the 120° F_1_ power strokes (8). Based on this asymmetry observed in low resolution ADP-inhibited F_1_F_o_ structures (26), high efficiency TD formation was proposed to occur (27) in the rotary state comparable to that in which rotary sub-state structures PDB-IDs 6OQR and 6OQS were subsequently observed at 3.1 Å resolution (17). Sobti *et al*. (17) concurred that torsional strain contributed to their ability to resolve the 6OQR and 6OQS sub-state structures. However, in results presented here, catalytically active F_1_F_0_ in lipid bilayer nanodiscs show successive 11° ATP synthase steps every 36° including at the rotary position of the ATPase power stroke where ADP inhibits rotation (**Fig 2A**, 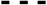). Because ATP synthase steps can also occur with low efficiency when torsional strain decreases the probability of forming a synthase step, it is clear that torsional strain is not the primary contributing factor to the ability of F_o_ to undergo 11° ATP synthase steps.

The data presented here are consistent with a Grotthuss proton translocation mechanism through the two subunit-a half channels connected by the proton transfer events to and from the c-ring. In a Grotthuss mechanism (28), a chain of water molecules that is H^+^-bonded to specific protein groups, enables transfer of protonic charge over long distances via rapid exchange of H^+^between H_3_O and H_2_O. This type of proton transfer is supported, first, by the fact that the mutations of all residues investigated here caused significant changes in the ability to form TDs, including the pH dependence of TD formation, and the ability to form 11° CW synthase steps in particular. This indicates that these groups all participate in the H^+^ transfer process. In addition, none of the mutations completely eliminated the pH dependent 11° ATP synthase steps, consistent with the requirement that a Grotthuss-type water column must be supported by multiple H-bond partners to enable H^+^ translocation (28,29).

Second, in structure 6OQS (17), the putatively protonated cD61-carboxyl comes within 3.5 Å of the polar aS199-hydroxyl, which is incapable of accepting a proton by itself. However, the 4.6 Å distance between aS199 and aE196 is consistent with the presence of an intervening water such that deprotonation of cD61 would be able to generate a hydronium ion that would immediately protonate the aE196-carboxyl. Similarly, residues aN214 and aQ252 at the input channel:c-ring interface are not ionizable, and thus are unable to directly protonate cD61. However, these polar sidechains may protonate cD61 if their role is to provide H-bonds to a Grotthuss water column. The other input channel residues aH245, aE119, and aE219 are all separated by distances of 4 Å to 7 Å that can stabilize a Grotthuss water column. It is noteworthy that the path between these residues includes backbone carbonyls that can also contribute to the stabilization of a H^+^translocating water column. Although a potential path for the output channel between aE196, and the cytoplasm has not been identified, aQ181, aE177, and the subunit-a C-terminal carboxyl group span this distance at ~4 Å intervals (17), consistent with that needed to stabilize a Grotthuss water channel in the *E. coli* enzyme. However, more work is required to characterize this channel, especially since aE196 and aS199 are the only output channel residues conserved among other species.

Third, the data presented here show that the high and low pKa values correspond to the H^+^-transfer events from the input channel to the c-ring, and from the c-ring to the output channel, respectively. However, mutation of a residue in either half-channel alters both pKa’s. The good fit of the occurrence of ATP synthase steps with the proportion of groups with high and low pKa values in the correct protonation state, supports the conclusion that synthase steps occur when H^+^-transfer occurs between the c-ring and both half channels. Conversely, H^+^-transfer events between the c-ring one of the half channels result in TDs that lack a synthase step. As the result of an 11° CW synthase step, a periplasmic proton entering the input channel Grotthuss water column would result in the immediate release of a proton from the output channel Grotthuss column into the cytoplasm. It is noteworthy that ATPase-driven CCW rotation pumps protons in the opposite direction from that which occurs during ATP synthesis. Although this process is reversible, the results presented here that showed that ATP synthase steps increased with aQ252L and especially aN214L, decreased the efficiency of H^+^-transfer in the ATPase direction relative to the ATP synthase direction.

Fourth, H^+^-specific conductance through FO from *Rhodobacter capsulatus* was observed to increase linearly with the size of a transmembrane voltage jump from 7 to 70 mV (30). A conductance of ~10 fS, equivalent to a proton translocation rate of 6240 H^+^ s^−1^ was observed at 100 mV driving force. Such high rates of H^+^ translocation support a Grotthuss mechanism, and are so fast that the ability to supply protons to the Grotthuss water column becomes a rate-limiting factor (29). The rate clearly exceeds the rate of delivery of protons by free diffusion from the bulk aqueous solution at a concentration of 10^−8^ M (pH 8). To achieve this rate of H^+^ translocation, the existence of a proton antenna at the distal end of the F_o_ input channel has been postulated (29), which in *R. capsulatus* was calculated to consist of a hemispherical Coulomb cage with a H^+^capture radius of ~40 Å surrounding the entrance to the input channel (29). The Coulomb cage would need to contain unprotonated carboxylate residues with a pKa ≅ 5. It is noteworthy that the pKa values estimated for *R. capsulatus* F_o_ (28) are comparable to those reported here for the groups that must be protonated to induce TDs.

In fact, *E. coli* F_1_F_o_ subunit-a (17) has a funnel shaped vestibule (**Fig 6**). The funnel diameter is ~30 Å at its widest as defined by surface polar groups, and is lined with several carboxylate and imidazole residues as the funnel narrows, culminating at its apex with the aE219-carboxyl examined here, which we propose to be at the start of the Grotthuss column. A recent cryo-EM structure of the V_O_ complex (31) was of sufficient resolution to resolve a column of water in the proton translocation channels of subunit-a of this related proton pumping rotary ATPase. Unidentified electron densities have also been identified near input channel residues in subunit-a in F_1_F_o_ structures from *E. coli* (17), and from *Polytomella* (32) that may indicate the presence of bound waters.

## Materials and Methods

### Mutagenesis and Purification of n-F_o_F_1_

The *E. coli* F_o_F_1_ samples were expressed from the pNY _1_-Ase plasmid construct with 6-His tag on the N-terminus of subunit-β, and a cysteine inserted at the second position of subunit-c (c2VCys) described previously by Ishmukhametov *et al*. (9). The aN214L, aQ252L, aH245L, aE219L, and aE196L point mutations were generated by site-directed mutagenesis. XL10-Gold Ultracompetent *E. coli* cells (Agilent) were transformed with the plasmid, the F_o_F_1_ complex was purified by detergent solubilization and Ni-NTA affinity chromatography, biotinylated, and incorporated into lipid nanodisc as previously described (8).

### Gold-Nanorod Single Molecule Experiments

Rotation of individual n-F_o_F_1_ molecules were observed by single-molecule rotation assay. Sample slides were prepared with modifications of previously described methods (8,9). Briefly, purified n-FOF1 were immobilized on a microscope slide by the His-tag on subunit-β, unbound enzymes were washed off the slide with wash buffer (30 mM Tris, 30mM PIPES, 10 mM KCl, at the appropriate pH), 80 × 40 nm AuNR coated with avidin was bound to the biotinylated c-ring of *E. coli* n-F_o_F_1_, excess AuNRs were washed off with the wash buffer, and rotation buffer (1 mM Mg^2+^ ATP, 30 mM Tris, 30mM PIPES, 10 mM KCl, at the pH indicated) was added to the slide. The rotation of individual molecules was observed by measuring the change in intensity of polarized red light scattered from the AuNR using a single-photon detector. In each molecule observed, the rotation of the nanorod attached to an active n-F_o_F_1_ complex was confirmed by the change in the dynamic range of the scattered light intensity as a function of the rotational positions of the polarizing filter as described previously (18,19). To make the measurement of n-F_o_F_1_ undergoing power strokes, the orientation of the polarizing filter was adjusted to align with the minimum light intensity position that that corresponded to one of the three catalytic dwells. The sinusoidal change of polarized red light intensity was measured as the AuNR rotated from 0° to 90° relative to the catalytic dwell position. Measurements were taken in the form of 5 s dataset at frame rate of 100 kHz. The occurrence of transient dwells in each subunit-a mutant was analyzed at varying pH from 5.0 to 8.0. Transient dwells that occurred during the power strokes in the recorded data sets were analyzed using custom software developed in MATLAB R2103b (10, 33).

## Acknowledgments

This work was funded by NIH grant R01GM097510 to WDF.

## Competing Interests

The authors declare that they have no competing interests. All data needed to evaluate the conclusions in the paper are present in the paper and/or the Supplementary Materials.

## Supplementary Materials

**Fig. S1.**
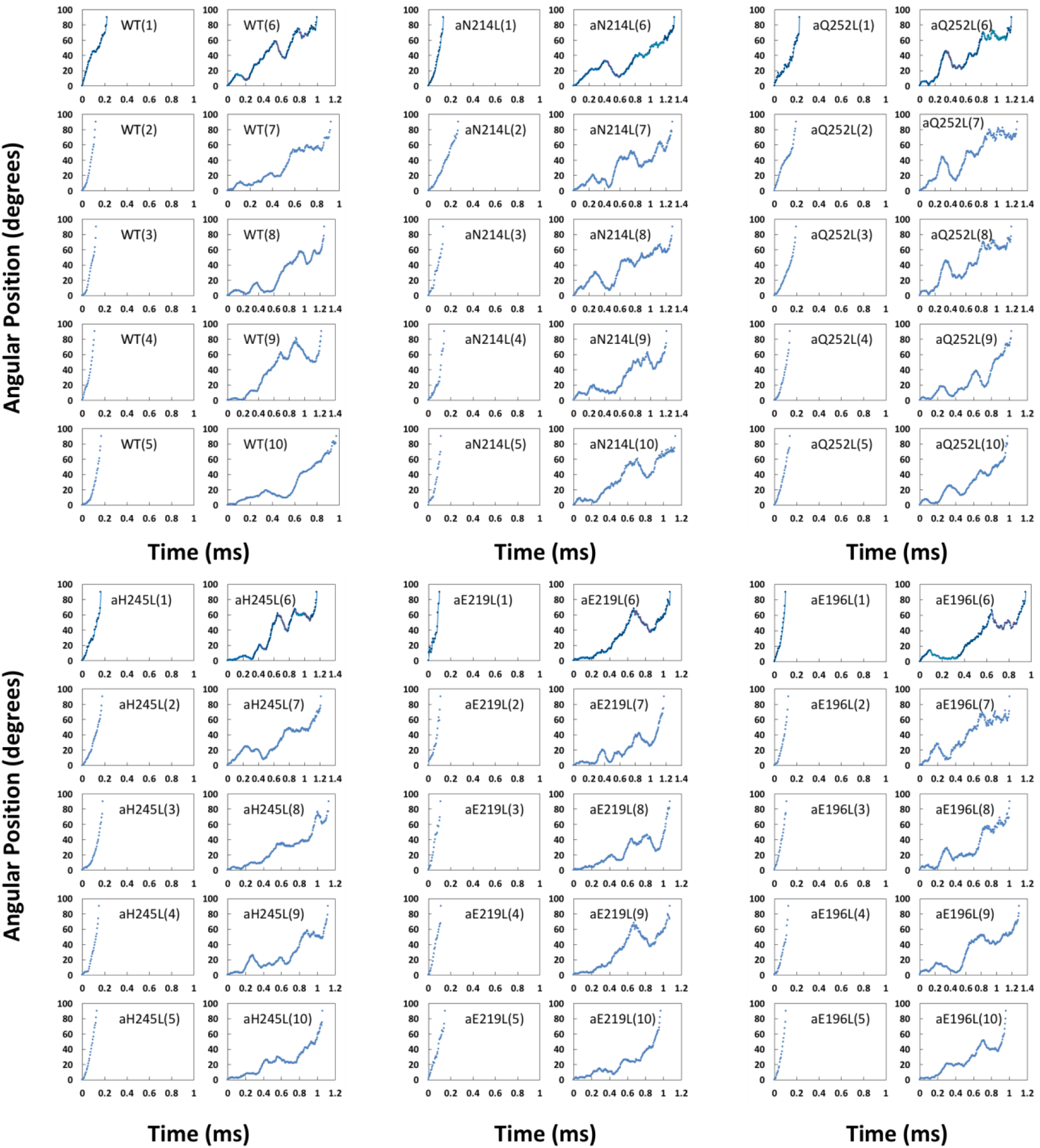
Examples each of the first 90° of ATP hydrolysis-driven power strokes observed using F_o_F_1_-nanodiscs. In each mutant, examples 1-5 show power strokes without transient dwells. Examples 6-10 show power strokes with transient dwells where the F_o_ motor either halts CCW rotation, or caused CW rotation in the ATP synthase direction.

**Fig. S2.**
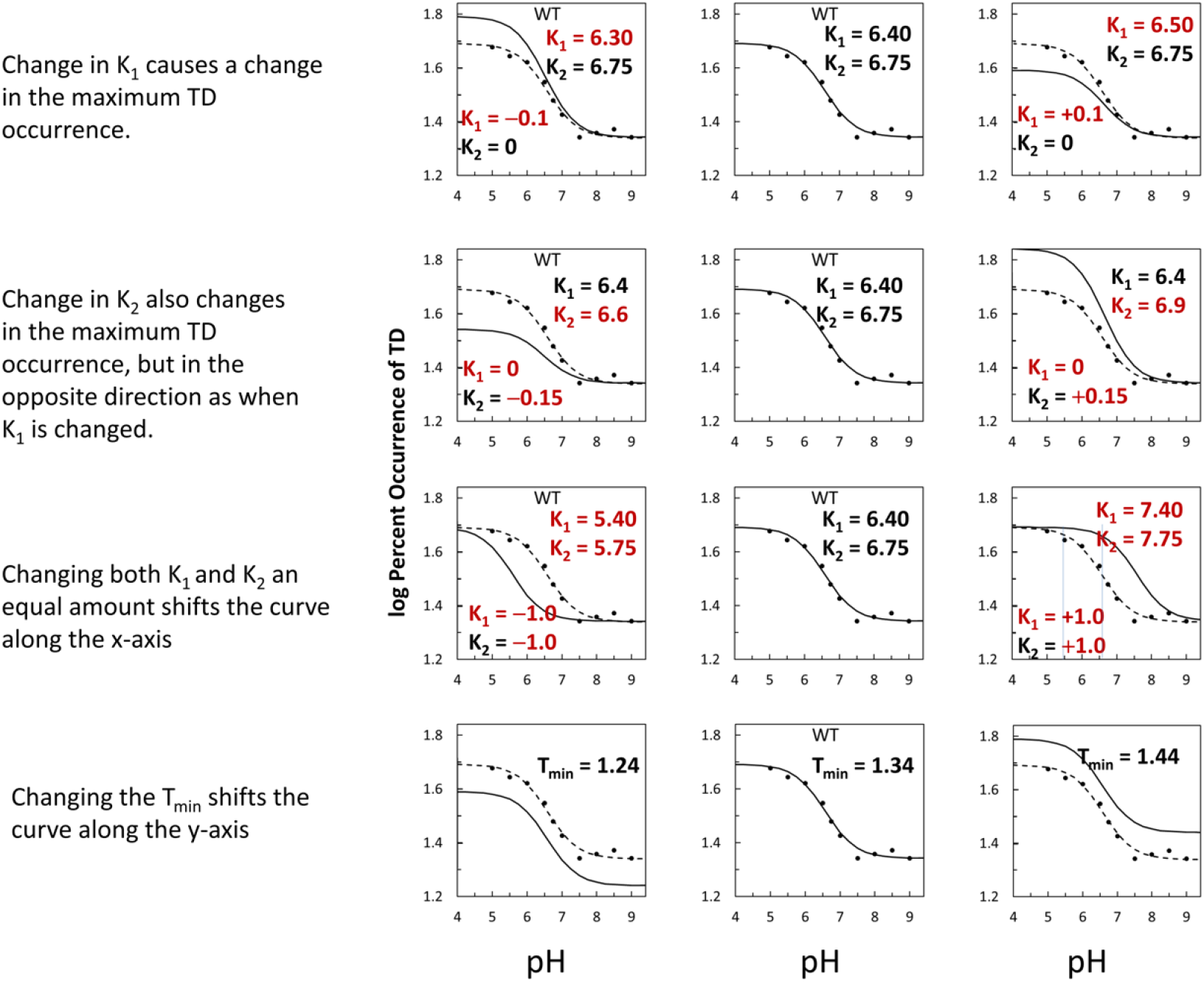
Examples of how changes in the variables in *Eq. 1* affect the log-log plots that describe the F_1_-ATPase inhibition kinetics of Fig 2C.

**Fig. S3.**
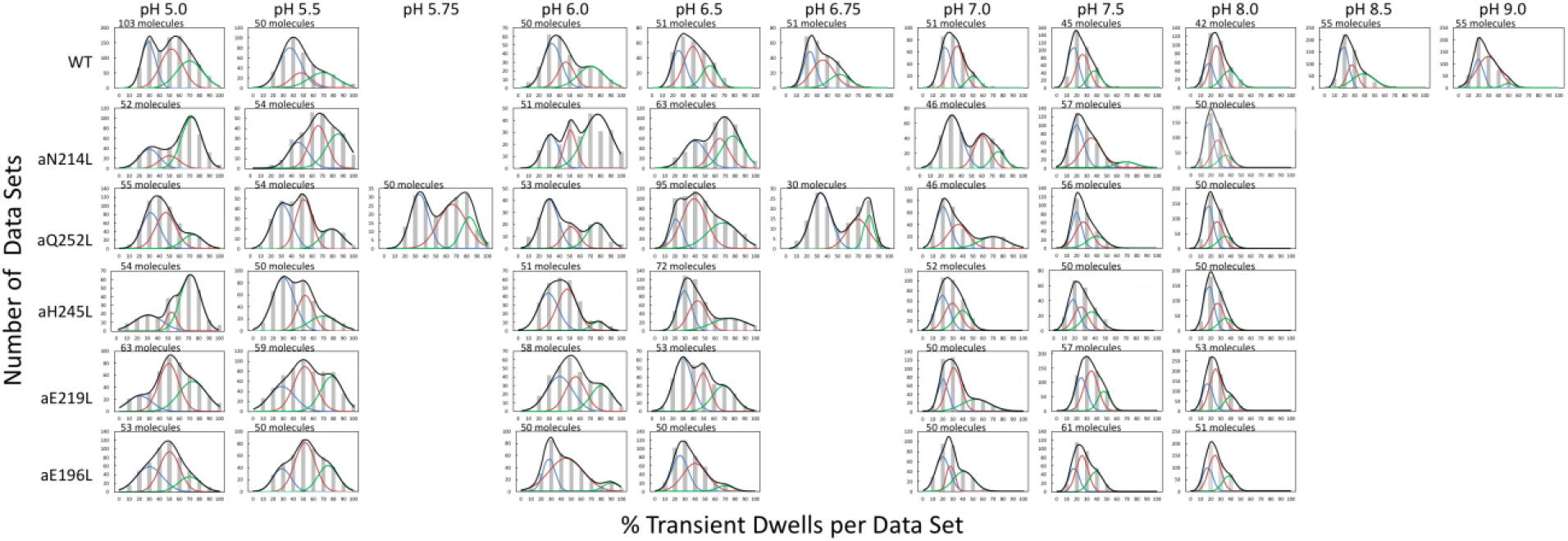
Distribution of power stroke data sets (each set containing ~300 power strokes) at each pH examined *vs* the percentage of the occurrence of TDs per data set binned to each 10 % (gray bars). The data were fit to the sum of three Gaussians 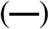 representing low 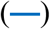, medium 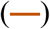, and high 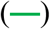 efficiencies of TD formation. Ten data sets were acquired from each molecule, and the number of molecules examined are shown for each condition.

**Fig. S4.**
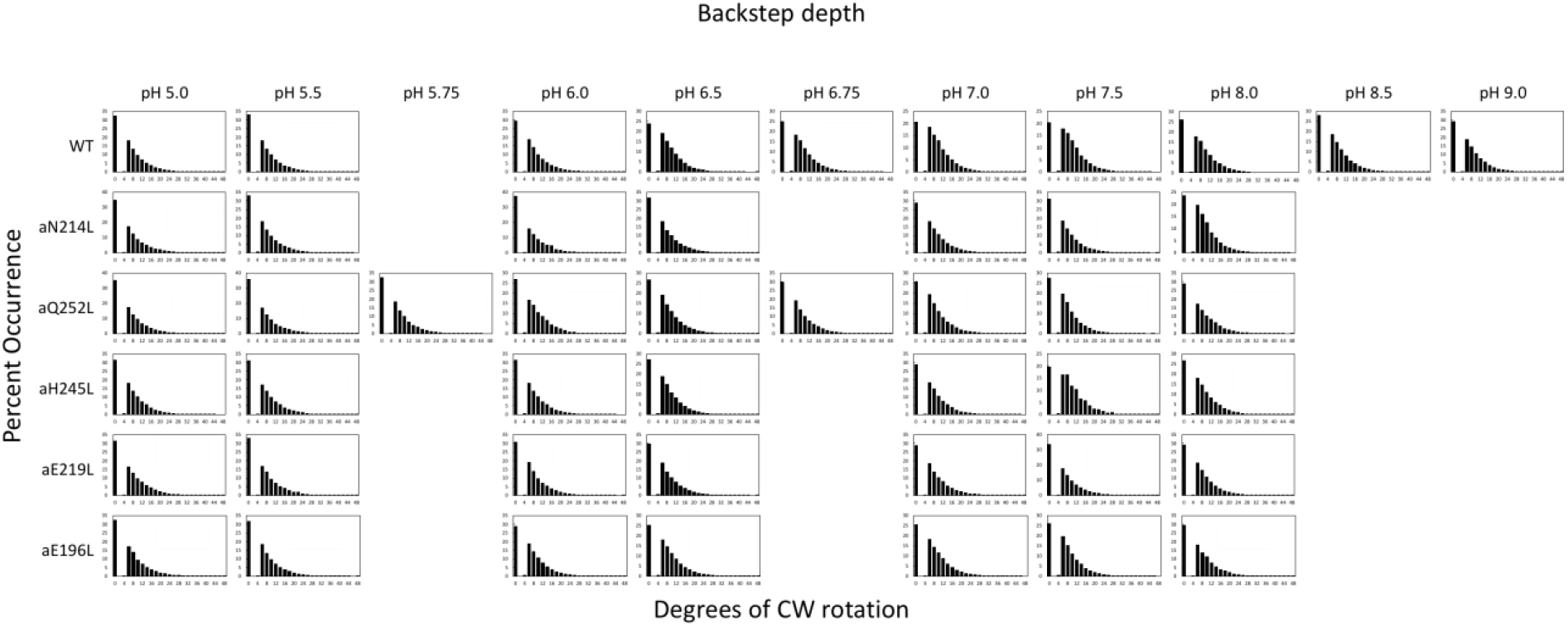
Distributions of the extent of CW rotation in the ATP synthesis direction during transient dwells for WT and subunit-a mutants *vs* pH.

**Fig. S5.**
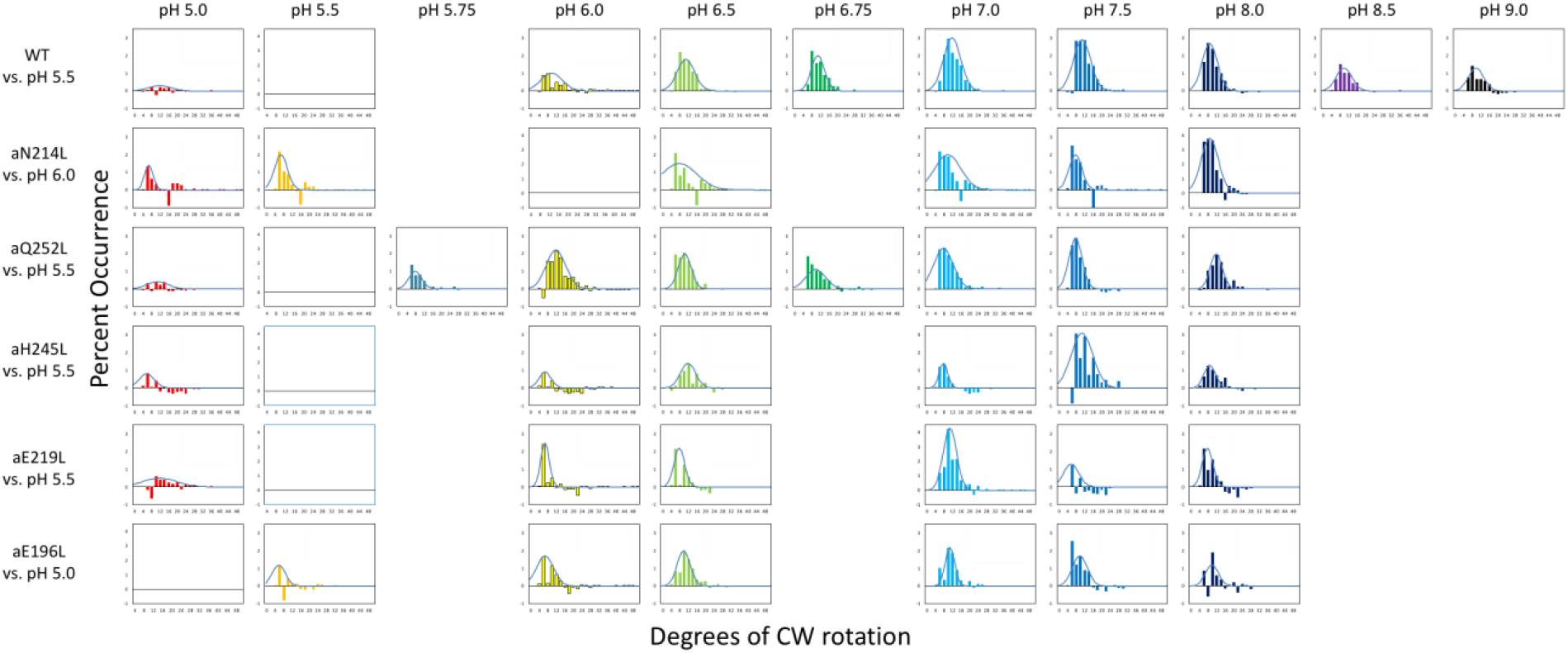
Distributions of the difference in extent of CW synthase step rotation between pH values when the percent of synthase steps was maximum *vs* minimum, where 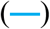 is the Gaussian fit.

## Notes

### Competing Interest Statement

The authors have declared no competing interest.

